# ProTrack: An Interactive Multi-Omics Data Browser for Proteogenomic Studies

**DOI:** 10.1101/2020.02.05.935650

**Authors:** Anna Pamela Calinawan, Xiaoyu Song, Jiayi Ji, Saravana Mohan Dhanasekaran, Francesca Petralia, Pei Wang, Boris Reva

## Abstract

The Clinical Proteomic Tumor Analysis Consortium (CPTAC) initiative has generated extensive multi-omics data resources of deep proteogenomic profiles for multiple cancer types. To enable the broader community of biological and medical researchers to intuitively query, explore, and download data and analysis results from various CPTAC projects, we built a prototype user-friendly web application called “ProTrack” with the CPTAC clear cell renal cell carcinoma (ccRCC) data set (http://ccrcc.cptac-data-view.org). Here we describe the salient features of this application which provides a dynamic, comprehensive, and granular visualization of the rich proteogenomic data.

**Statement of Significance:** The CPTAC initiative (https://proteomics.cancer.gov/) has generated multi-omics data for multiple cancer types to understand the proteogenomic aberrations of these malignancies. Collectively this effort has so far produced a large data resource for the research community, including high-throughput profiles for proteome, phosphoproteome, whole exome, whole genome, transcriptome, and DNA methylome. To make this valuable data-resource useful to the larger research community, there is a pressing need for development of user-friendly, readily accessible, and easily shared analytic and visualization tools for aligning multi-omics data and exploring alterations in key cancer genes, to drive and support new biological hypotheses. To bridge this gap, we have developed CPTAC ProTrack, an interactive web application which uses a multilayered, client-server architecture in order to deliver an interactive web experience to any user of a modern web-browser. This tool is intentionally designed accessible for researchers, biologists, and clinicians who are interested in multi-omic data without any need to code.

---

For example, screenshots of the browser-based user interface shown in **Figure 1** helps identify somatic mutation, CNV and immune infiltration related markers associated to different immune clusters. Clearly, the tool helps visualize the enrichment of patients with BAP1 mutation and Chr14 deletion in CD8+ Inflamed group (Fig 1A). In addition, it helps to display key immune cells markers and checkpoint inhibitors based on multi-omic data. As shown, CD8A is elevated in the CD8+ immune group; while VEGFA in the VEGF immune desert group. Immune checkpoint inhibitors PD-1 (PDCD1), PD-L1 (CD274) are clearly elevated in CD8+ inflamed group. Therefore, ProTrack can help identifying patterns of association between multi-omic data with a particular phenotype of interest such as immune clusters and clinical information. In addition, through ProTrack users can efficiently generate visual heatmaps of customized combinations of multi-omic data from CPTAC and perform an initial query of such associations. In this paper, we focus on introducing ProTrack for the CPTAC clear cell renal cell carcinoma (ccRCC) study [1], importantly similar tools for CPTAC proteogenomic studies of other cancer types, will be made available as and when the corresponding data is released by the consortium or upon publication.

**Fig. 1.**
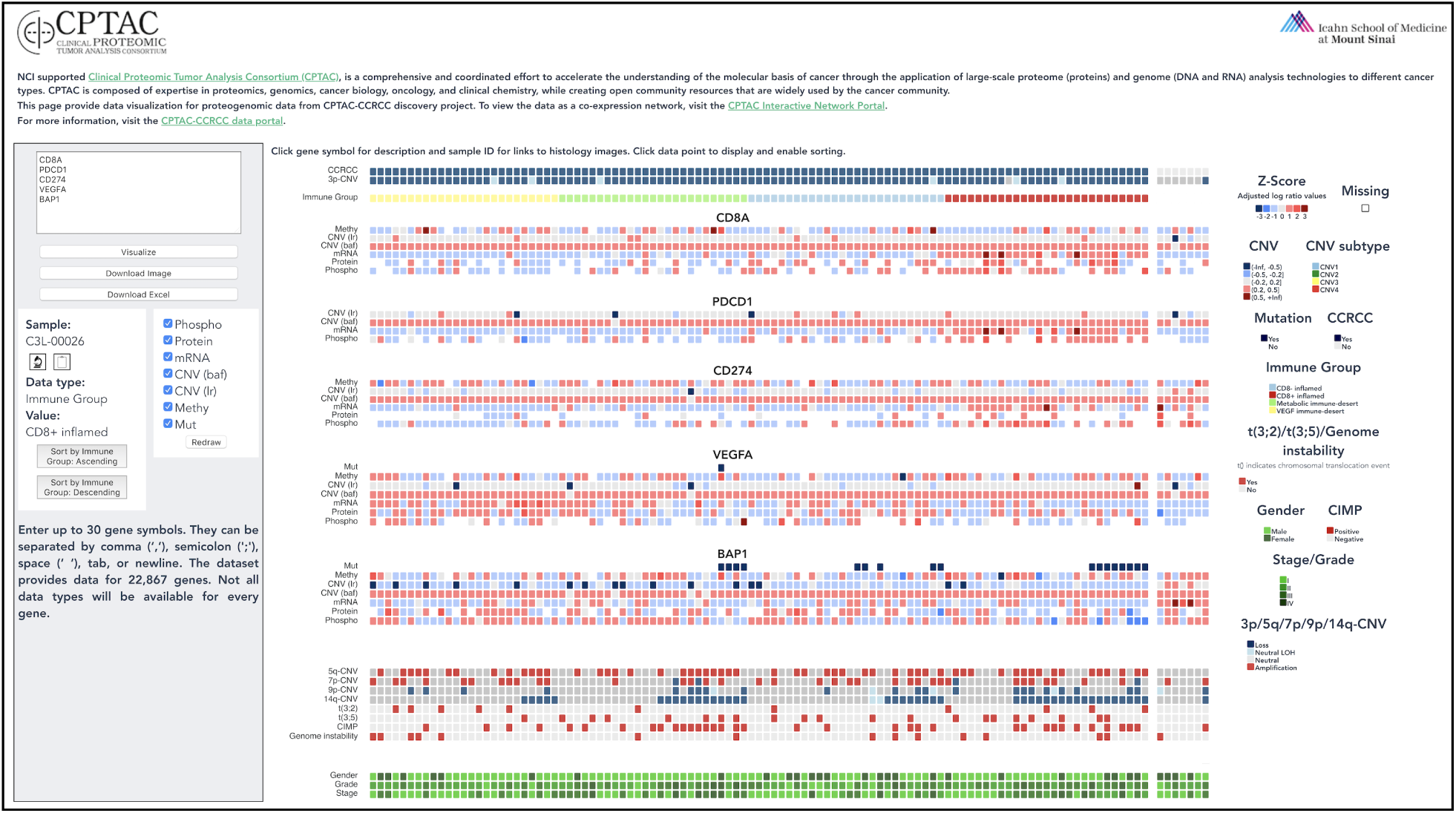
Screenshot of ProTrack. Samples are sorted based on the four immune clusters identified in ccRCC: CD8+ Inflamed Group (red), CD8- Inflamed Group (lightblue), Metabolic Immune Desert (green) and VEGF Immune Desert (yellow). Multi omic data such as mutation profiles, gene expression and protein expression for CD8A, PD-1 (PDCD1), PD-L1 (CD274), VEGFA and BAP1 are shown along with CNV events and clinical information such as grade and stage.

ProTrack utilizes heatmap visualizations to facilitate multi-omics data viewing as they allow for alignment and visual inference across different data types, wherein each data type can be represented by a single track, and multiple omics layers can be organized per gene. Adjacent tracks for different omics layers allow users to immediately visualize data relationships for a single gene, across multiple genes, for a single sample, or across all samples as required. In contrast to a static heatmap image, users can also inspect the values underlying an individual data point. In addition, ProTrack contains interactive functionality which allow users to select data types such as proteomics, phosphoproteomics and RNAseq data to be visualized for a given set of genes, view sample identifiers, and explore patterns by efficiently sorting samples based on either clinical or molecular data. This multi-omics visualization tool will enable researchers to immediately explore new findings and facilitate comparisons of findings between cancer data cohorts.

## ccRCC Data Description

Renal cell carcinoma (RCC), one of the top ten most commonly diagnosed cancers worldwide, is dominated by the clear cell histological subtype, which makes up 75% of all RCC cases [1]. To understand the molecular alterations underlying the ccRCC disease process, CPTAC has performed an extensive multi-omic characterization of 110 treatment-naive tumors and 84 paired normal adjacent tissues samples. In addition to clinical annotations, the underlying big-data resource includes molecular annotations for 11,355 proteins and 42,889 phosphopeptides, as well as whole genome sequencing (WGS), whole exome sequencing (WES), RNA sequencing (RNA-Seq), and methylation data. ProTrack organizes this data into top and bottom tracks of genetic and clinical annotations, including genomically confirmed ccRCC, non-ccRCC status; 3p copy number variation; immune subtype of each sample; copy number variation for chromosomes 5q, 7p, 14q; chromosomes 2,3 and 3,5 translocations; genome instability; CpG island methylator phenotype (CIMP) status; grade; stage; and gender. Gene-level data can be represented by multiple tracks, when available, including mutation status (“Mut”), the average beta value of methylation probes in the CpG island of in the promoter region of the gene (“Methy”), the log ratio of copy number variation (“CNV (lr)”), the b-allele frequency of the copy number variation (“CNV (baf)”), gene expression levels (“mRNA”), gene-level protein abundance (“proteo”), and gene-level phosphoprotein abundance (“phospho”). Full details on the statistical summarization of the underlying data can be obtained from our recent publication (ref: Clark et al).

## Comparison to Existing Tools

There are some existing limitations for the tools that currently exist tofor viewing and exploringe multi-omic cancer data. In the field of interactive heatmaps, portals such as Next-Generation Clustered Heat Map (NG-CHM) Viewer [2] and iCoMut Beta for FireBrowse [3] readily exist for exploring and generating heatmaps of molecular cancer data. But tThese portals are specifically designed to host data from utilize pre-populated data from The Cancer Genome Atlas (TCGA), which focused on genomic, epigenomic, and transcriptomic profiling with limited proteomic data.

There are also broader service platforms that allow users to upload custom data sets and render interactive heatmaps, such as PaintOmics3 [4] and Clustergrammer [5]. The limitation of these generic and powerful platforms is, when researchers intend to collaborate with a common data set,. uUploading data from CPTAC, or another data source, can beis time-consuming and lead to data duplication. In addition, and manipulation investigation of the data by from one researcher to another in ways that can be especially difficult to trace and reproduce by another researcher. Furthermore, grappling with a one-size-fits-all visualization tool can be quite complex and requires all users to be experienced with this specific tool. With a simple, dedicated tool that is coupled to a fixed underlying dataset, collaborators can reproduce, explore, and expand upon each other’s data visualization with minimal need for data transport among collaborators. Moreover, the single-purpose feature set has a much lower learning curve to produce visually pleasing and useful output for data exploration.

Accordingly, ProTrack has been comprehensively designed as an easy-to-learn service for users interested in these specific CPTAC datasets. ProTrack allows the researcher to access a centralized source of data rather than bringing one of many copies of data to the tool, facilitating consistent and reproducible analyses.

While the OncoPrint tool on cBioPortal [6] includes CPTAC data and a host of advanced features for data analysis and visualization, its multi-omic heatmap visualization is, at the time of writing, still limited to protein level Z-scores for the available CPTAC tumor, while ProTrack also includes phosphoproteomic and methylome tracks. In general, cBioPortal offers a much more extensive and advanced array of cancer analysis and visualization tools, but requires much more effort, expertise, and refinement to produce meaningfully targeted output. The simplicity of ProTrack allows users to explore the data set without specific expertise for this platform. Simply by inputting a gene set, researchers can get immediately started with exploring data without grappling with an overwhelming array of options. In its most basic use-case, ProTrack is input-output: enter a list of genes and receive a heatmap.

## Feature Workflow

**Figure 2A** shows a workflow diagram for the visualization interface of ProTrack. First the user submits up to 30 genes of interest and an interactive heatmap is rendered in their browser. Once the visualization is rendered, the user can interact at the node and track level. At the node level, users can click an individual node to see the sample identifier and underlying data value. ProTrack accesses external web services through their application programming interfaces (APIs) to act as a one-stop shop for CPTAC data. When a node is selected, the user can see histologic images from the Cancer Imaging Archive; gene descriptions from the National Center for Biotechnology Information; and demographic annotations from the Proteomic Data Commons. At the track level, users can toggle tracks on and off by selecting the data types of interest and clicking ‘Redraw.’ Gene-level tracks are labeled with the gene symbol, which the users can click to read a description of the gene from the National Center for Biotechnology Information (NCBI). Clicking a track enables the sorting feature. Users can sort a selected track in ascending or descending order, and the entire heatmap will rearrange according to the sample order determined by the sorted track.

**Fig 2.**
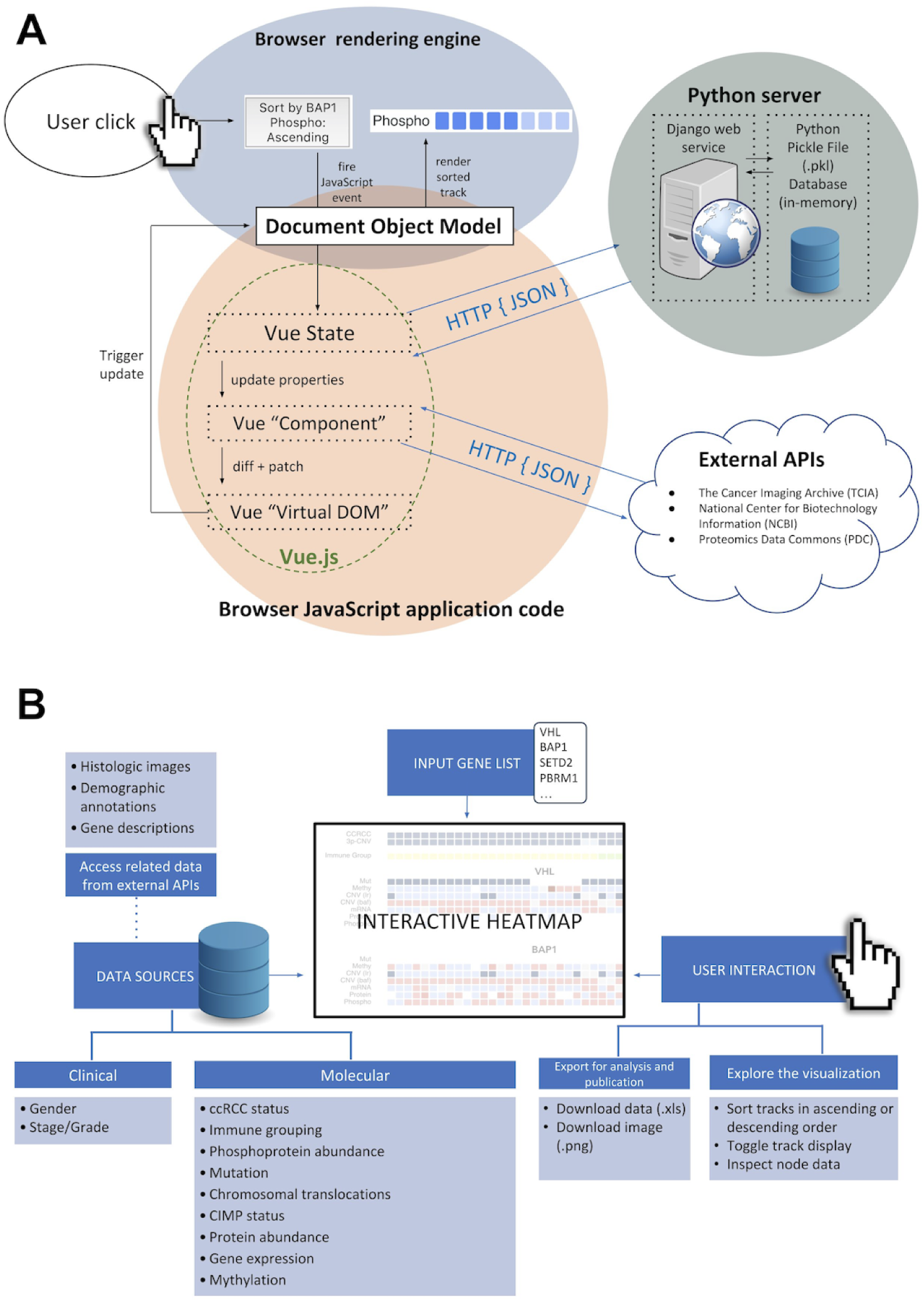
A) A simplified schema of the data flow for a single user action and its flow through the application. In this example, a user clicks a button intending to render a sorted heatmap track. User actions trigger JavaScript functions, as built within the Vue framework. Data transformations occur within the application, while necessary changes to the visual template are calculated by Vue in the “Virtual DOM” layer. This virtual DOM selectively manipulates visual elements on the page, and renders them accordingly. Moreover, the Vue data layer fetches data, when required, from the Python backend with an in-memory static data store. Additional CPTAC data is dynamically fetched from external data sources. B) A summary of the data and interactive features available in ProTrack.

Users can also download the image for future reference and export the underlying data table as an Excel file, sorted in exactly the same order that produced the heatmap rendered at the time of clicking “Download Excel”.

The amount of data that ProTrack can display at one time is limited by various factors. The user’s computer resolution and screen size can limit how effectively the visualization can be perceived and explored. The speed of the user’s Internet connection and the computing power of the system determine how quickly the heatmap will render.

## Software Design and Implementation

This software uses a multilayered, client-server architecture in order to deliver an interactive web experience to any user of a modern web-browser. This multilayer architecture, illustrated in **Figure 2B**, has the advantage of specializing and segregating implementation technologies between the server and client, using technologies best-suited to their distinct purposes in the broader application.

The server (or “backend”) uses the web framework Django. Django is a popular Python-based web framework that emphasizes speed, security, and scalability for web backends. It handles the various important tasks related to forming a response to HyperText Transfer Protocol (HTTP) requests, including request parsing, path routing, data serialization, and composing a standardized message responses. In ProTrack, the Django backend acts as a RESTful API layer, providing response data in a standardized and optimized JavaScript Object Notation (JSON) format for client consumption.

Moreover, by using a Python-based backend, the web application can easily integrate with Python’s enormous availability of data processing libraries that are strongly optimized for efficient data processing tasks written in a high-level programming language. Accordingly, pre-processed data are stored in-memory in the running Django python task as Python Pickle files. Because the stored data to be delivered to the client is read-only, this effectively acts as the entire backend database layer for the application. On the client-side (or “frontend”), this project leverages the cutting-edge of frontend web technologies in order to provide a fast user experience for exploring, visualizing, and sharing CPTAC data. ProTrack is built on modern web standards that have been adopted by all modern browsers. Vue, a JavaScript library and web-development framework, is used as the primary library to build the client-side application. By contrast to traditional interactive webpages which combine statically rendered HTML documents with selective layers of interactivity through JavaScript, ProTrack has a single-page application (SPA) architecture where all elements are generated via JavaScript, and all user interactions dynamically transform the rendered content directly versus loading a completely new page. Vue aids in the development of client-side SPAs by using the Virtual DOM, a data tree model of the rendered page content, which smartly optimizes webpage rendering when small portions of visual information change based on user input. This application was written with Vue’s “single file component” file structure, further streamlining the development process by combining Hypertext Markup Language (HTML) templating, JavaScript, and Cascading Style Sheets (CSS) visual styling in a single file. Additionally, JavaScript tooling such as Webpack and Babel, aid cross-browser compatibility and delivering to the client optimized JavaScript code bundles, increasing user download and execution speeds. Together with the Vue library, the application also uses ApexCharts as its visualization library, which emphasizes performance with large data sets and a flexible API for a catered visualization experience.

## Visualizing mutually exclusive translocation events in ccRCC

The value of the heatmap interactivity can be demonstrated in the following example. In the ccRCC study by Clark et al, the novel chromosome 3-2 rearrangement, consisting primarily of 3p loss and 2q gain, was observed to be nearly mutually exclusive to 3-5 translocation events. This mutual exclusivity is readily demonstrated by sorting the interactive heatmap by t(3;2) in descending order followed by t(3;5) in descending order, as shown in **Figure 3**.

**Figure 3.**
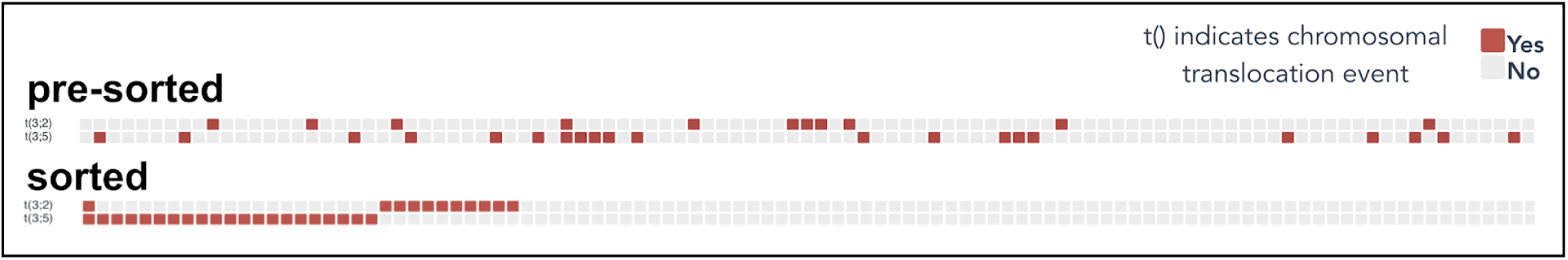
The t(3;2) and t(3;5) translocation events shown here are nearly mutually exclusive, which is not readily shown by the unsorted heatmap. Sorting the heatmap in descending order by t(3;2) followed by descending order by t(3;5) readily demonstrates this mutual exclusivity.

## Data and Software Access

The tool is entirely open-source and source code and pickle files containing the underlying data are available on Github at http://www.github.com/WangLab/ProTrack-ccrcc. The hosted data can also be downloaded as Excel files from the website at http://ccrcc.cptac-data-view.org.

## Conclusion

We have leveraged recent developments in web technologies to create an intuitive and interactive tool for granular, customized exploration of multi-omic data, starting with the ccRCC cohort, including proteomic data that is not readily available on any other visualization platform. CPTAC proteogenomic studies of other cancer types will include discovery and confirmatory cohorts, and as this data is released by the consortium or published, we will continue to develop interactive heatmaps customized to each data type. The interactive data and visualizations will be aggregated into a single platform to facilitate analysis across multiple cancer types.

## Acknowledgments

This work was supported by the NIH, National Cancer Institute’s Clinical Proteomic Tumor Analysis Consortium (CPTAC) grants U24CA210985, U24CA210993, U24CA210967, U24CA210954, and U24CA210972.

## Abbreviations

CPTAC, ccRCC, WGS, WES, RNA-Seq, TCGA, NCBI, HTML, CSS, CIMP, NCBI, PDC, TCIA

